# Proximity proteomics reveals the molecular architecture of phytochrome B photobodies in *Arabidopsis thaliana*

**DOI:** 10.1101/2025.11.09.687487

**Authors:** Kate Olson, Jae-Hyung Lee, Madison Dean, Thu Minh Doan, Allison M. Manuel, Chan Yul Yoo

## Abstract

The red/far-red light photoreceptor phytochrome B (phyB) forms light-induced subnuclear condensates, termed photobodies, that coordinate plant responses to light and temperature. Despite their central role in environmental signaling, the molecular composition of phyB photobodies during their early formation has remained unknown. Here, we established an *in planta* proximity labeling approach using miniTurbo-based biotinylation in *Arabidopsis thaliana* to capture proteins associated with phyB photobodies during early de-etiolation. Mass spectrometry identified 42 high-confidence proximal proteins, including 11 known core components and 31 previously unrecognized photobody-associated proteins. Among these, the co-chaperone HOP1 forms light-dependent nuclear condensates that partially co-localize with phyB photobodies. HOP1 condensates are smaller in the wild-type background than in phyB-overexpressing seedlings, and HOP1 overexpression enhanced cotyledon expansion under red light. These findings suggest that HOP1 contributes to photomorphogenesis by stabilizing phyB photobodies and sustaining active phyB signaling. Together, our results reveal that nascent large phyB photobodies function as dynamic hubs integrating chaperone-mediated protein quality control with transcriptional regulation, providing the first *in planta* proteomic framework for understanding photobody assembly and signaling in plants.

## INTRODUCTION

Plants continuously monitor fluctuating environments to optimize growth, metabolism, and survival. Light is not only a source of energy but a fundamental environmental signal that shapes seedling photomorphogenesis (Von Arnim & Deng, 1996). During germination in darkness, etiolated seedlings display elongated hypocotyls, tightly closed cotyledons, and undeveloped plastids. Upon exposure to light, seedlings rapidly undergo de-etiolation, a developmental reprogramming characterized by hypocotyl growth inhibition, cotyledon opening and expansion, chloroplast biogenesis, and activation of photomorphogenic gene expression (Chen *et al*., 2004; Kami *et al*., 2010). In *Arabidopsis thaliana*, the red/far-red light photoreceptor phytochrome B (phyB) is a master integrator of light and temperature cues, functioning as both a photoreceptor and thermosensor (Rockwell *et al*., 2006; Jung *et al*., 2016; Legris *et al*., 2016; Cheng *et al*., 2021). Upon red light activation, phyB shifts from the inactive cytoplasmic Pr form to the active Pfr form, translocates into the nucleus, and assembles into subnuclear condensates called photobodies that initiate photomorphogenesis (Yamaguchi *et al*., 1999; Kircher *et al*., 1999; Van Buskirk *et al*., 2012; Zhao *et al*., 2023).

Photobodies are light-induced biomolecular condensates whose formation depends on multivalent interactions and intrinsically disordered regions (IDRs) (Willige *et al*., 2024). The N-terminal extension (NTE) of phyB contains a critical IDR required for photobody assembly, and NTE-deficient phyB mutants fail to rescue photomorphogenic defects (Chen *et al*., 2022). Photobodies act as hubs that coordinate two major light-signaling pathways: suppression of the COP1/SPA ubiquitin ligase complex and degradation and inhibition of PHYTOCHROME-INTERACTING FACTORS (PIFs) (Chen, 2008; Chang *et al*., 2025). COP1/SPA actively represses photomorphogenesis in darkness by destabilizing positive transcriptional regulators such as HY5. Red light-activated phyB inhibits COP1/SPA activity and stabilizes light-responsive transcription factors (Zheng *et al*., 2013; Sheerin *et al*., 2015). In parallel, photoactivated phyB interacts with PIF1, PIF3, PIF4, PIF5, and PIF7 for phosphorylation and ubiquitin-mediated degradation, and inactivation of their transcriptional activities, allowing transcriptional reprogramming of photosynthetic and developmental programs (Bauer *et al*., 2004; Willige *et al*., 2021; Kim *et al*., 2023a). Core photobody scaffolds such as PHOTOPERIODIC CONTROL OF HYPOCOTYL 1 (PCH1) and PCH1-LIKE (PCHL) stabilize the photoactivated Pfr state and promote formation of large photobodies (Huang *et al*., 2016, 2019; Enderle *et al*., 2017). In addition, the photobody-localized regulator TANDEM ZINC-FINGER PLUS3 (TZP) fine-tunes hypocotyl elongation by regulating phyB protein abundance and photobody formation, and PIF signaling (Kaiserli *et al*., 2015; Fang *et al*., 2022), underscoring photobodies as hubs that couple light perception to developmental output. Thus, photobodies function as dynamic condensates that integrate protein turnover, transcriptional control, and environmental signaling.

A major advance toward defining photobody composition came from a recent proteomic studies that used fluorescence-activated particle sorting (FAPS) to isolate photobody-enriched nuclear fractions from *phyB-GFP* plants, followed by liquid chromatography-tandem mass spectrometry (LC-MS/MS) (Kim *et al*., 2023b). However, because FAPS-based isolation involves mechanical disruption of nuclei, it likely recovers photobody-containing particles rather than intact photobodies and may not capture transient or weak associations, distinguish photobodies formed under different environmental conditions, or profile early photobody formation during de-etiolation.

To overcome these limitations, we developed an *in planta* proximity labeling strategy using miniTurbo-based biotinylation to identify proteins associated with nascent large phyB photobodies during early de-etiolation. This approach captures protein interactions in living cells at nanometer resolution, enabling discovery of transient, low-abundance, or unstable photobody components (Branon *et al*., 2018; Qin *et al*., 2021). Our analysis revealed 42 high-confidence proximal proteins, including all three HEAT SHOCK PROTEIN 70-HEAT SHOCK PROTEIN 90 ORGANIZING PROTEINS (HOP1, HOP2, and HOP3), a family of conserved co-chaperones that mediate client transfer between HSP70 and HSP90 complexes for regulating protein stability. We further showed that HOP1 formed light-dependent condensates that partially co-localized with phyB photobodies. HOP1 overexpression enhanced cotyledon expansion under red light. These results suggest that HOP1 acts as a co-chaperone that stabilized large photobodies to maintain active phyB signaling. Together, these findings establish the first *in planta* proteomic framework for understanding the molecular architecture and functional dynamics of phyB photobodies.

## RESULTS

### Establishing an *in planta* proximity labeling system to identify phyB photobody components

To enable direct *in planta* identification of proteins associated with phytochrome B (phyB) photobodies, we established a proximity labeling strategy using the engineered biotin ligase miniTurbo (Fig. 1a). To implement this system, we generated Arabidopsis lines expressing phyB fused to miniTurbo, cyan fluorescent protein (CFP), and a C-terminal FLAG tag under the *UBIQUITIN10* (*UBQ10*) promoter in the *phyA-211/phyB-9* (*phyA/phyB*) double mutant background (PB-mTb) (Fig 1b). As a negative control, we introduced a previously published nuclear miniTurbo-yellow fluorescent protein (YFP)-nuclear localization signal (NLS) construct (mTb) into the same bcaackground (Mair *et al*., 2019). Two independent PBmTb lines (#3-1 and #10-1) fully rescued the elongated-hypocotyl phenotype of *phyA/phyB* seedlings under red light, while the mTb control failed to complement the mutant phenotype (Fig. 1c, d). PBmTb also restored cotyledon expansion and markedly increased chlorophyll accumulation relative to the *phyA/phyB* mutant background (Fig. 1c, e).

**Figure 1.**
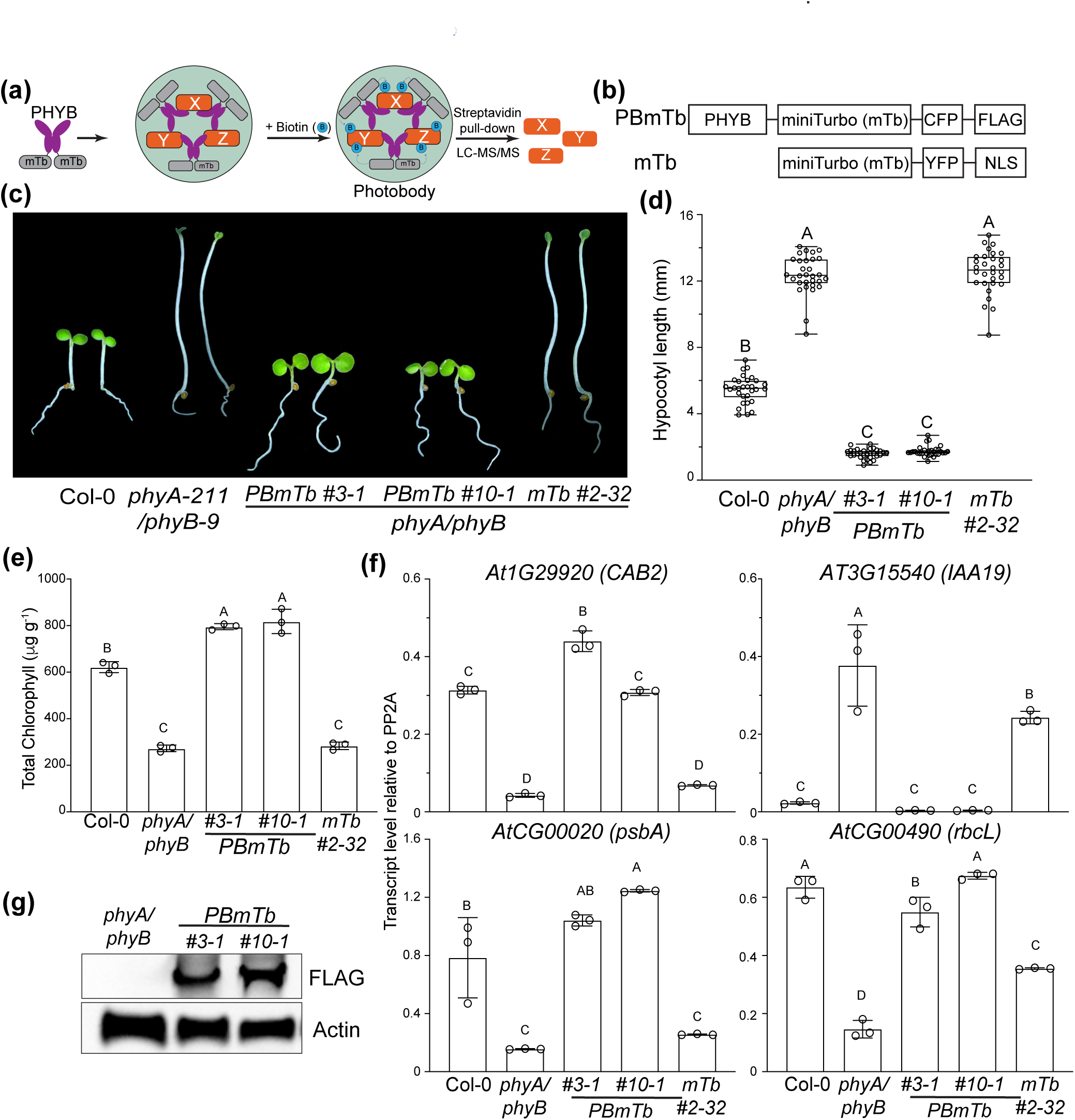
Functional validation of phyB-miniTurbo for proximity proteomics in Arabidopsis. (a) Schematic illustration of phyB-miniTurbo-mediated proximity labeling strategy to identify photobody-associated proteins. (b) Schematic illustration of the domain structures of PBmTb (phyB-miniTurbo-CFP-FLAG) and the nuclear control mTb (miniTurbo-YFP-NLS). (c) Representative images of 4-day-old Col-0, *phyA-211/phyB-9* (*phyA/phyB*), *PBmTb #3-1, PBmTb #10-1*, and *mTb #2-32* seedlings grown in 20 μmol m^-2^ s^-1^ R light. (d) Box-and-whisker plots showing hypocotyl lengths of seedlings shown in (c). Boxes indicate the 25^th^ to 75^th^ percentiles with median values shown as horizontal lines; whisker extends to the minimum and maximum values. (e) Total chlorophyll levels in seedlings shown in (c). (f) Transcript levels of phyB-regulated nuclear genes (*CAB2* and *IAA19*) and plastid-encoded photosynthetic genes (*psbA* and *rbcL*) in 4-d-old indicated lines grown in 20 μmol m^-2^ s^-1^ R light. (g) Immunoblot analysis of PB-mTb proteins detected with anti-FLAG antibody. Actin was used as a loading control. For (d), (e), (f). different letters indicate statistically significant differences (one-way ANOVA with Tukey’s HSD test, *P* < 0.01).

To evaluate restoration of phyB-mediated signaling for photomorphogenesis in the PBmTb lines, we analyzed representative nuclear and plastid genes involved in light-regulated development and chloroplast biogenesis (Yoo *et al*., 2019a). Expression of *CAB2* and IAA19, which respectively indicate red-light-induced chloroplast development and auxin-regulated hypocotyl growth inhibition, were fully recovered in *PBmTb* seedlings but not in the *mTb* control. Similarly, plastid-encoded *psbA* and *rbcL* transcripts were restored to wild-type levels in *PBmTb* lines (Fig. 1f), indicating proper activation of both nuclear and plastid branches of the phyB signaling network. Immunoblot analysis detected full-length phyB-miniTurbo fusion proteins in both lines, with *PBmTb #10-1* showing slightly higher levels than *#3-1* (Fig. 1g). Under continuous far-red light, *PBmTb* seedlings remained elongated like *phyA/phyB* mutants (Fig. S1a), confirming that phyB-miniTurbo does not interfere with phyA-mediated signaling. *PBmTb #10-1* seedlings exhibited slightly shorter hypocotyls than *#3-1* (Fig. S1b). Based on these molecular and phenotypic differences, we selected the *PBmTb #3-1* line for subsequent analyses. Together, these results demonstrate that miniTurbo tagging preserves phyB function and that the *PBmTb #3-1* line provides a reliable system to investigate the photobody dynamics and molecular composition of phyB photobodies.

### Light-dependent and photoreversible behavior of phyB-miniTurbo photobodies during de-etiolation

To verify that the phyB-miniTurbo fusion (PBmTb) faithfully reproduces photobody behavior, we examined PBmTb-CFP localization in dark-and light-grown seedlings. In darkness, PBmTb-CFP fluorescence was diffuse in the cytoplasm, whereas red-light exposure induced discrete photobody formation in the nucleus (Fig. 2a). In white-light-grown seedlings, PBmTb photobodies dispersed into a diffuse nuclear pattern when treated with far-red light, demonstrating light-dependent reversibility of photobody formation (Fig. 2b). During dark to light de-etiolation, numerous small nuclear foci appeared by 4 hour and progressively matured into larger photobodies by 24 hour (Fig. 2c). Quantification showed a time-dependent increase in both photobody volume and the number of large photobodies per nucleus, with values at 24 hour light exposure still smaller than under continuous red (R_C_) (Fig. 2d, e). These results demonstrate that PBmTb undergoes characteristic light-dependent assembly and disassembly cycles.

**Figure 2.**
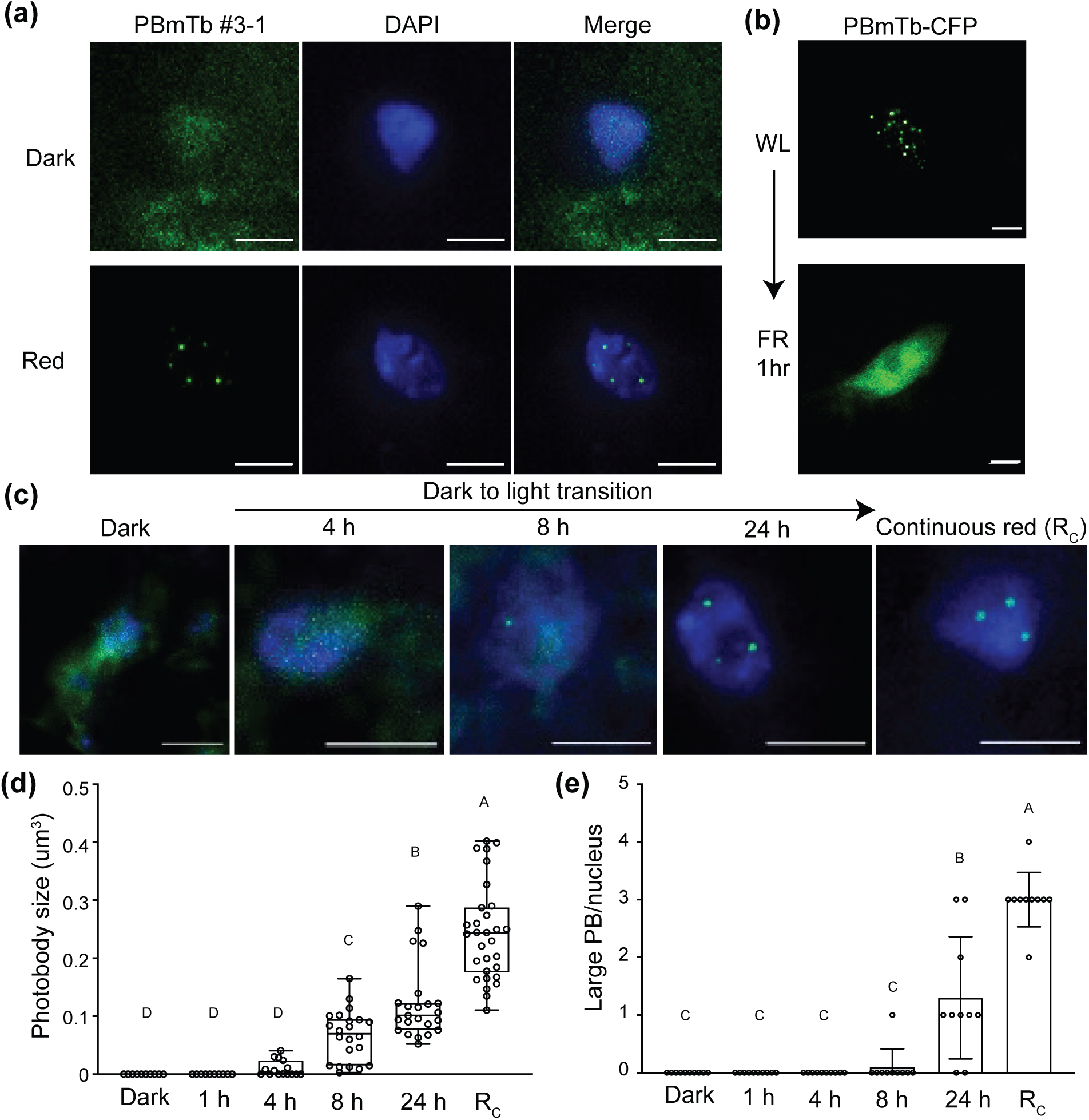
Light-dependent and photoreversible behavior of phyB-miniTurbo photobodies during de-etiolation. (a) Microscope images of 4-day-old PBmTb-CFP #3-1 seedlings grown in dark and 36 μmol m^-2^ s^-1^ red-light. (b) PBmTb-CFP localization in white-light-grown seedlings after 1 hour of far-red (FR) treatment. (c) Time-course images showing PBmTb-CFP photobody formation during dark-to-red light transition (1, 4, 8, and 24 hour) and in continuous red light (R_C_). (d) Quantification of mean photobody volume and (e) number of large photobodies per nucleus in seedlings shown in (c). Boxes indicate the 25^th^ and 75^th^ percentiles with median values shown as horizonal lines; whickers extend to the minimum and maximum values. Different letters indicate statistically significant differences (one-way ANOVA with Tukey’s HSD test, *P* < 0.01). Scale bars = 5 μm.

To validate that our experimental conditions corresponded to an early stage of de-etiolation, we compared seedling morphology and plastid gene expression under identical conditions (Fig. S2). After light exposure, seedlings exhibited only partial cotyledon expansion, and transcript levels of plastid-encoded photosynthetic genes (*psbA* and *rbcL*) increased relative to dark but remained below level from seedlings grown under continuous red light. Together, these results indicate that our light conditions define an intermediate stage of de-etiolation characterized by the emergence of large phyB photobodies. This condition was therefore used in subsequent experiments to examine early events in photobody maturation and composition.

### Proximity proteomics defines the early-stage phyB large photobody proteome

To determine the composition of phyB large photobodies during early de-etiolation, we performed proximity-dependent biotin labeling using 4-day-old PBmTb and mTb control seedlings. Dark-grown seedlings were exposed to 24 hours of red light to induce large photobodies, then submerged in 50 μM biotin for 3 hours under same light condition. Immunoblotting with Streptavidin-HRP confirmed robust labeling in PB-mTb lines, with the #10-1 line showing higher overall signal intensity consistent with its elevated protein level (Fig. 3a). The PBmTb #3-1 and mTb #2-32 control line were therefore used for subsequent affinity purification and mass spectrometry analyses. Label-free proteomic analysis of streptavidin-purified proteins identified over 5,000 proteins across three biological replicates (Fig. 3b). Principal component analysis (PCA) revealed clear separation between PBmTb and mTb samples, indicating strong experimental reproducibility and distinct enrichment profiles. (Fig. S3a). As expected, phyB was the most enriched protein in PBmTb samples, reflecting self-biotinylation of the fusion protein and confirming efficient miniTurbo activity and recovery of the bait protein (Fig. S3b).

**Figure 3.**
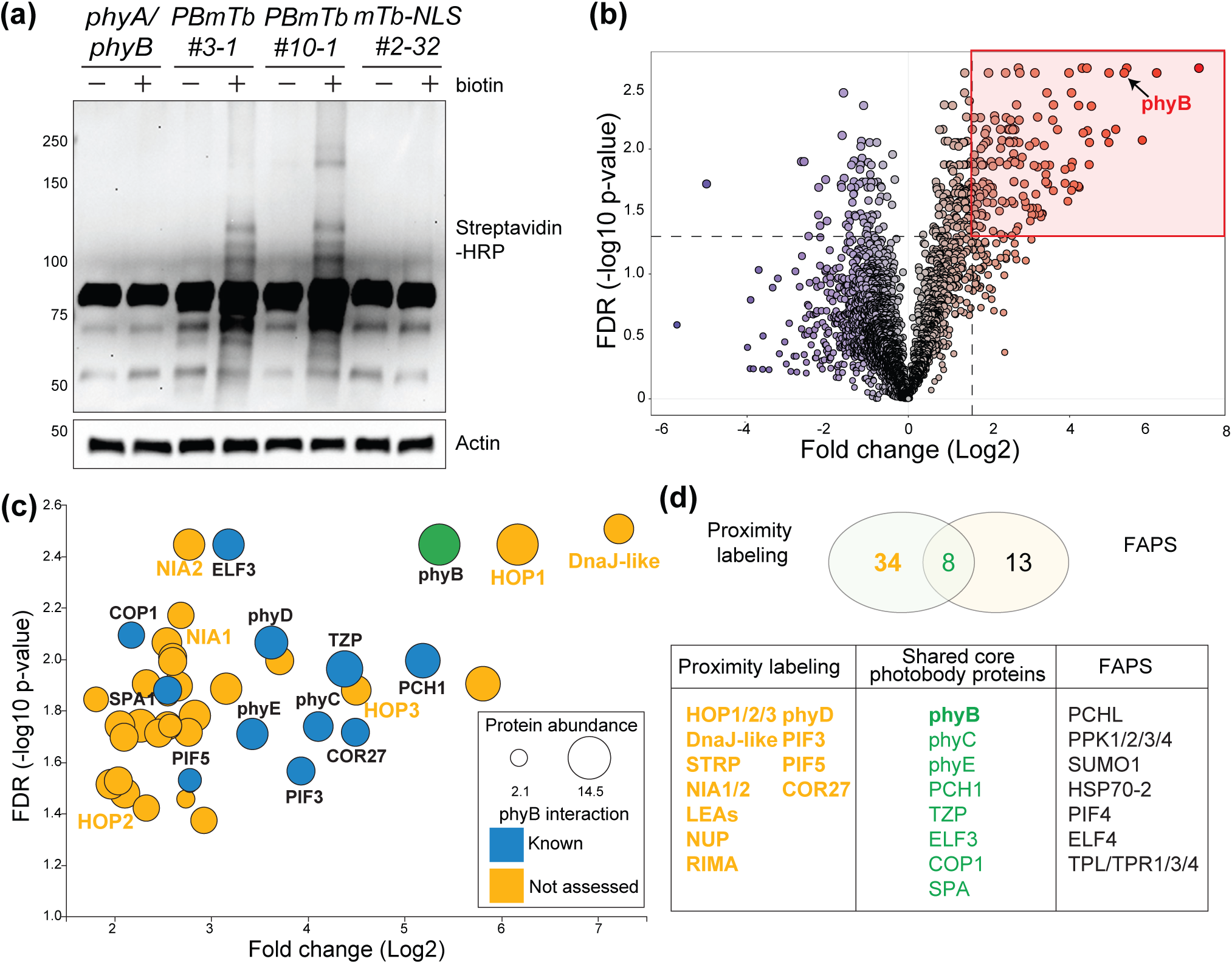
Proximity proteomics defines the early-stage phyB photobody proteome. (a) Immunoblot analysis of biotinylated proteins in 4-day-old *phyA/phyB, PBmTb* and *mTb* control seedlings exposed to 24 hour of 36 μmol m^-2^ s^-1^ red-light followed by 3 hour incubation in 50 μM biotin under light, detected with Streptavidin-HRP. Actin was used as a loading control (b) Volcano plot of proteins quantified in PBmTb #3-1 versus mTb control. Proteins above the threshold (fold change > 3 and FDR-adjusted *P* < 0.05) were considered significantly enriched and indicated in the red box. (c) Selected enriched proteins plotted by fold change (log2) and statistical significance. Circle size indicates protein abundance (log2); color indicates prior evidence of phyB interaction (blue, known; orange, not assessed). (d) Overlap between the PBmTb proximity-labeling dataset and the phyB photobody proteome obtained by fluorescence-assisted particle sorting (Kim *et al*., 2023b). Numbers in the Venn diagram denote proteins unique to each method and those shared; the table lists examples from each category.

We first applied moderate enrichment threshold (fold change > 3, FDR < 0.05), we identified 119 proteins enriched in PBmTb samples (Fig 3b, Table S1). We further removed proteins predicted to localize to organelles and visualized the distribution of enrichment significance, displaying proteins across a range of adjusted *P-*values, including the most stringent cutoff (*P* ≤ 0.01). This stringent included phyB and previously characterized phyB interactors including ELF3, COP1, phyD, PCH1, TZP, and SPA1, validating dataset accuracy (Fig. 3c, Table S2). In the moderate range (0.01 < FDR < 0.05), our data set also included additional known phyB interactors such as phyC, phyE, COR27, PIF3, and PIF5, further validating the reliability of the proximity labeling dataset for low abundance phyB interactors (Table S2). In addition to these known components, we detected a suite of previously uncharacterized proteins potentially associated with early large photobodies, including molecular chaperones (HOP1/2/3, DnaJ-like HSP40), stress-related protein STRP, nitrate reductase (NIA1/2), intrinsically disordered proteins (e.g. LEA proteins), and several nuclear envelope-associated proteins (NUP) (Fig. 3c). The presence of these factors suggests that early large photobodies may integrate protein quality control and stress-response modules as well as mediators of nucleocytoplasmic communication.

To assess conservation of photobody components, we compared our dataset with a recently reported phyB photobody proteome obtained by FAPS (Kim *et al*., 2023b). Eight proteins were shared between the two datasets, representing conserved core photobody components (Fig. 3d). Several secondary interacting proteins such as PPK family kinases, HSP70-2, and TPL, which have been identified as indirect phyB partners through PCH1 or PIF3, were not detected in our dataset. Conversely, our proximity labeling uniquely identified 34 additional proteins, including known phyB interactors such as phyD, PIF3, PIF5, and COR27,which may represent transient or weakly associated factors enriched in early photobodies during de-etiolation. Collectively, these analyses define a comprehensive early-stage phyB photobody proteome and reveal new candidate regulators that may contribute to photobody assembly and environmental responsiveness.

### HOP1 forms light-dependent condensates associated with phyB photobodies and promotes cotyledon expansion

Among the newly identified photobody-associated proteins, the co-chaperone HOP1 was of particular interest because all three Arabidopsis HOP family members (HOP1, HOP2, HOP3) were recovered in the photobody proteome (Fig. 3d). To examine whether HOP1 associates with phyB photobodies, YFP-HOP1 was expressed under the *UBQ10* promoter in *phyB-CFP* (PBC) and Col-0 backgrounds. As a positive photobody marker control, YFP-TZP was also expressed in PBC background. In red-light grown seedlings, YFP-TZP co-localized with phyB-CFP exclusively at photobodies, as expected. YFP-HOP1, by contrast, formed discrete condensates in both the nucleus and cytoplasm, with a subset of nuclear foci overlapping with phyB photobodies (Fig. 4a).

**Figure 4.**
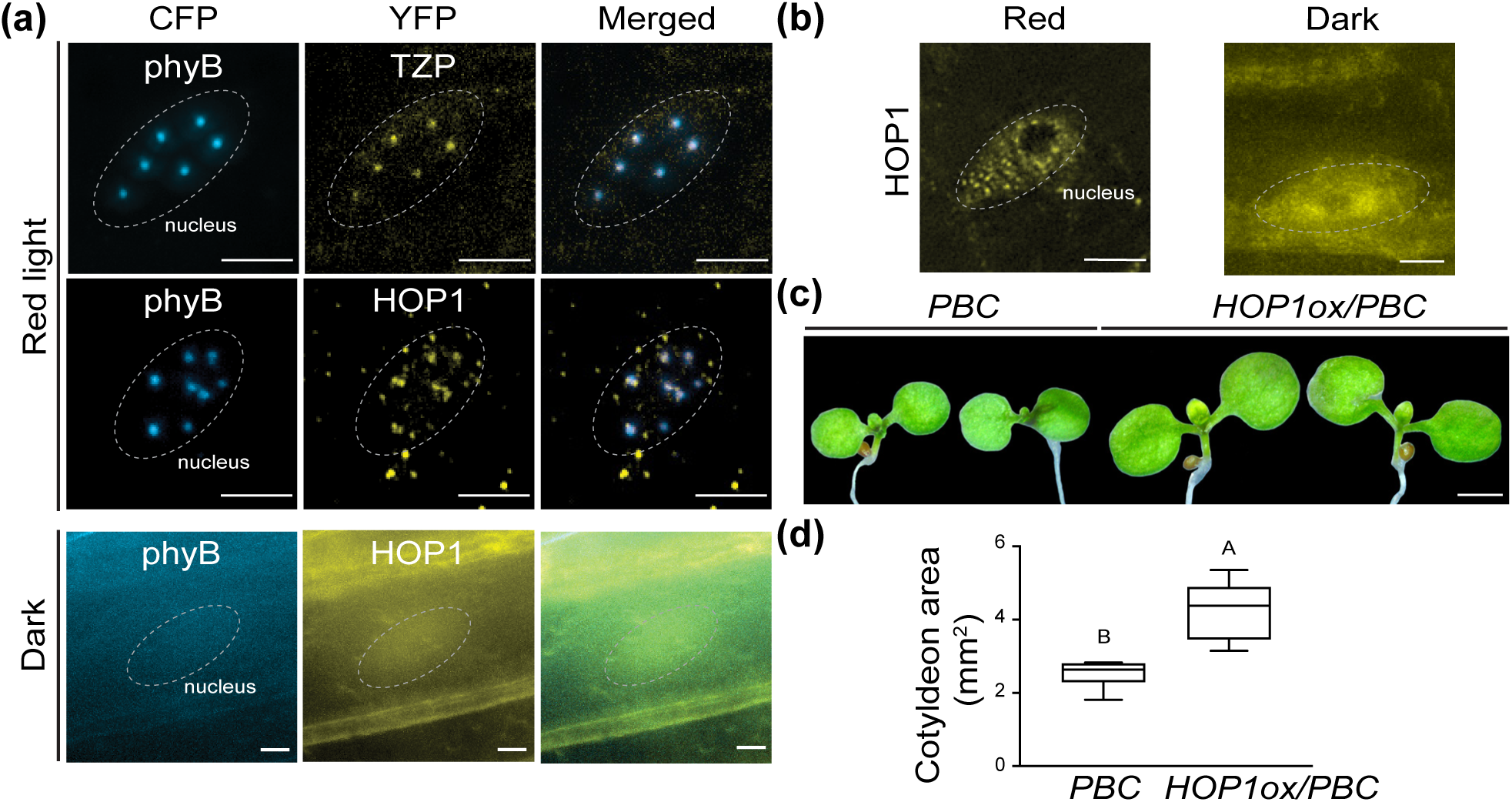
HOP1 forms light-dependent condensates associated with phyB photobodies and promotes cotyledon expansion. (a) Microscope images of YFP-HOP1 and YFP-TZP expressed under the *UBQ10* promoter in *phyB-CFP* (*PBC*) background. Seedlings were grown in darkness or under continuous 20 μmol m^-^_2_ s^-1^ red-light. Scale bars = 5 μm. (b) YFP-HOP1 localization in Col-0 background under dark and red-light conditions. Scale bars = 5 μm. (c) Representative images of 4-day-old red-light-grown *PBC* and *YFP-HOP1/PBC* seedlings. Scale bar = 1 mm. (d) Box-and-whisker plots showing cotyledon area of seedlings shown in (c). Boxes indicate the 25^th^ and 75^th^ percentiles with median values shown as horizonal lines; whickers extend to the minimum and maximum values. Different letters indicate statistically significant differences (Student’s *t* test, *P* < 0.01).

In darkness, YFP-HOP1 displayed diffuse localization in both compartments, indicating that HOP1 condensation is light-dependent (Fig. 4a). When YFP-HOP1 was expressed in the Col-0 background, where native phyB levels are lower, nuclear HOP1 condensates were still detectable but smaller in size compared to the PBC background (Fig 4b), indicating that phyB abundance modulates HOP1 condensation. These observations indicate that HOP1 undergoes light-induced phase transition and partially associates with phyB photobodies.

To assess the functional consequence of HOP1 in phyB signaling, cotyledon expansion was measured in YFP-HOP1 overexpression lines in PBC background and showed significantly enhanced size and area compared with PBC along (Fig. 4c, e). Together, these results demonstrate that HOP1 forms light-dependent condensates associated with, but not restricted to, phyB photobodies, The enrichment of HOP1 and related chaperones suggests that chaperone condensates may contribute to maintaining large phyB photobodies and sustaining active light signaling.

## DISCUSSION

Photobodies are hallmark subnuclear condensates that couple light perception with transcriptional and developmental reprogramming. Although photobody function is well established, the mechanisms governing their assembly and remodeling in living tissues remain poorly understood. Here, we developed an *in planta* miniTurbo-based proximity labeling approach to capture the composition of large phyB photobodies during their early formation in de-etiolating Arabidopsis seedlings. This strategy provides a physiologically relevant snapshot of protein associations surrounding phyB under native conditions.

Our proximity proteomics identified 42 high-confidence proteins enriched near phyB, including many canonical photobody components such as PCH1, TZP, ELF3, COP1, SPA, and PIFs, confirming the reliability of this approach. These proteins overlap extensively with those reported by Kim et al. (2023), who employed FAPS to isolate large photobodies from mature plants and identify the first photobody proteome by mass spectrometry. The overlap between these independent datasets supports the existence of a conserved core photobody module. However, our *in planta* labeling also revealed distinct features not observed in the FAPS-derived proteome. In particular, we found an enrichment of molecular chaperones, including all three HOPs (HOP1, HOP2, and HOP3) and DnaJ-like HSP40 protein. We also discovered various intrinsically disordered proteins including LATE EMBRYOGENESIS ABUNDANT (LEA) proteins that are abundant in seed and accumulated under stress including cellular dehydration tolerance. These disordered proteins might provide functional role in photobody formation. These unique proteins likely reflect the developmental stage and intact-cell context of our *in planta* labeling approach, which detects early, transient, and weak associations that may not exist in mature plants or may be lost during mechanical isolation of photobody-enriched fractions. Interestingly, we did not identify PIF4, but identify PIF3 and PIF5, suggesting that PIF3 and PIF5 play more functional role in photomorphogenesis. In addition, PIF4 protein levels are diurnally regulated and generally low at ZT 0 in our experimental condition. This might reflect different PIFs are associated with photobodies in different developmental and circadian stage. Together, these findings suggest that photobody assembly is accompanied by dynamic recruitment of chaperone and intrinsically disordered modules that support condensate nucleation and stability.

The identification of HOP1/2/3 highlights a previously unrecognized link between the chaperone network and light signaling condensates. HOP proteins act as co-chaperones bridging HSP70 and HSP90 to facilitate client protein folding, and they have been implicated in diverse signaling including thermomorphogenesis, thermotolerance, and other stress-response pathways (Fernández-Bautista *et al*., 2018; Toribio *et al*., 2020; Mangano *et al*., 2023). Surprisingly, HOP1 displayed light-dependent nuclear condensation that partially overlapped with phyB photobodies. HOP1 condensates were smaller in wild-type seedlings than in phyB-overexpressing backgrounds, indicating that phyB abundance influences HOP1 recruitment and condensate size. Interestingly, HOP1 remains diffuse in both the nucleus and cytoplasm of Col-0 and phyB-overexpressing lines in darkness, when inactive phyB is primarily localized in the cytoplasm. Future studies should test whether HOP1 condensates and phyB photobodies are interdependent. Moreover, overexpression of HOP1 enhanced cotyledon expansion under red light, suggesting that HOP1 positively promotes photomorphogenesis, possibly by stabilizing large photobodies or maintaining the active Pfr state of phyB. These findings, together with previous reports that HOPs contribute to thermomorphogenesis by stabilizing TIR1 auxin coreceptor (Muñoz *et al*., 2022) highlight a broader role for HOP co-chaperones in integrating light and temperature signaling through protein homeostasis.

The regulation of phyB photobody assembly likely parallels general principles governing chaperone-mediated control of biomolecular condensates. Recent studies in animal systems have shown that molecular chaperones not only prevent aberrant phase transitions but also actively remodel and disperse condensates under stress conditions (Yoo *et al*., 2022; Bard & Drummond, 2024). Canonical chaperones such as Hsp70 and small HSPs dynamically modulate condensate material properties by transiently binding disordered regions, thereby tuning the balance between condensation and dissolution. Similarly, engineered or non-canonical chaperones have been proposed to act as specificity factors that selectively regulate client condensates (Bushman *et al*., 2025).

Our results, when considered alongside the FAPS-based photobody proteome, provide a complementary perspective on the organization and dynamics of phyB condensates. The FAPS study revealed a mature photobody architecture composed of structural scaffolds and secondary interactors, whereas our *in planta* approach illuminates the early, chaperone-assisted phase of photobody formation in young seedlings. We propose a model in which phyB activation triggers the condensation of a chaperone-rich microenvironment that facilitates folding, protection, or exchange of client proteins during the transition from small puncta to stable large photobodies. As photobodies mature, structural and transcriptional regulators such as PCH1, TZP, and ELF3 become incorporated to execute stable signaling output.

Together, this study establishes the first *in planta* proteomic framework for phyB photobody assembly and reveals that chaperone-mediated quality control is an integral feature of light-dependent condensate biogenesis. These insights extend the current view of photobody function beyond static signaling compartments to dynamic proteostasis hubs that integrate environmental sensing, protein folding, and developmental regulation.

## MATERIALS AND METHODS

### Plant material and growth conditions

The *phyB-CFP* (*PBC*) line in the Col-0 background has been described previously (Kim *et al*., 2023b) and was used to characterize photobody behavior. The *phyA-211/phyB-9* double mutant (Reed *et al*., 1994) served as the background for constructs generated in this study. Additional Arabidopsis transgenic lines are described below. Seeds were surface-sterilized with 70% ethanol and bleach, then plated on half-strength Murashige and Skoog (MS) medium supplemented with Gamborg’s vitamins (MSP06, Casson Laboratories), 0.5mM MES pH 5.7, and 0.8% agar (w/v). Seeds were stratified at 4°C in the dark for 4 days. Seedlings were grown at 21°C under the indicated light conditions in an LED chambers (Percival Scientific). Light fluence rates were measured with a LI-180 spectrometer (LI-COR Biosciences). Representative seedling images were captured using a stereomicroscope and processed in Adobe Photoshop.

### Plasmid construction and generation of transgenic plants

To generate the *PBmTb* construct, *miniTurbo*, *SCFP3A* and *FLAG* were fused in the indicated order to the 3’ end of the phyB coding sequence driven by the *UBQ10* promoter. Fragments were amplified by PCR and assembled into the EcoRI and BamHI site of the pJHA212 vector using HiFi DNA assembly (New England Biolabs) (Yoo *et al*., 2005). The mTb control construct was obtained from Dominique Bergmann through Addgene (#127370) (Mair *et al*., 2019). Transgenic lines expressing *PBmTB* and *mTb* were generated by *Agrobacterium tumefaciens* (GV3101)- mediated transformation of *phyA-211/phyB-9* (Clough & Bent, 1998). Transformants were selected on MS medium containing 15μg mL^-1^ Basta, and multiple independent lines were established for analysis. To generate YFP-TZP and YFP-HOP1 constructs, *eYFP* and *3xHA* tag were fused to the 5’ end of *TZP* or *HOP1* coding sequences under the *UBQ10* promoter. Fragments were assembled into the XmaI and PstI sites of pJHA212 vector using HiFi DNA Assembly. Wild-type (Col-0) and *PBC* plants were transformed with *A. tumefaciens* carrying the respective constructs, and Basta-resistant lines were selected for subsequent experiments. Primers used to generate the plasmid constructs are listed in Table S3.

### Hypocotyl measurement

Seedlings were grown for 4 days under the indicated light conditions. At least 30 seedlings per genotype were scanned and analyzed for hypocotyl length with ImageJ software (NIH). Box-and whisker plots were generated using Prism 10 software (GraphPad).

### Chlorophyll measurement

Total chlorophyll was extracted from 100 mg of seedlings from each genotype in 3 ml of 100% dimethyl sulfoxide (DMSO) by incubation at 37°C for 30 min. Absorbance at 648.2 nm and 664.9 nm was measured by spectrophotometry, and total chlorophyll concentrations was calculated as described (Barnes *et al*., 1992).

### RNA extraction and quantitative reverse transcription PCR

Total RNA was extracted from 100 mg of seedlings of the indicated genotypes using the Quick-RNA MiniPrep Kit with on-column DNase I treatment (Zymo Research). First-strand cDNA was synthesized from 1 μg of total RNA using the Superscript IV cDNA Synthesis system according to the manufacturer’s protocol (Thermo Fisher Scientific). Oligo(dT) primers were used for nuclear gene expression analysis, a gene-specific primers were used for plastid transcripts. Quantitative RT-PCR was performed using the FastStart Essential DNA Green Master mix on a LightCycler 96 System (Roche). Transcript abundance of target genes were normalized to *PP2A* expression. Primer sequences used for cDNA synthesis and qRT-PCR are listed in Table S3.

### Protein extraction and immunoblot analysis

Total protein was extracted from *Arabidopsis* seedlings grown for 3 days in darkness and transferred to 36 μmol m^−2^ s^−1^ red light for 24 hours before flash freezing. Biotin-treated samples were prepared as described below. Approximately 100 mg of tissue was ground in liquid nitrogen and homogenized in extraction buffer (500 mM Tris-HCl pH 7.5, 150 mM NaCl, 0.1% SDS, 1% Triton X-100, 0.5% sodium deoxycholate, 1mM EGTA, 1mM DTT, protease inhibitor cocktail, 1mM PMSF, 40 μM MG132, 40 μM MG115). Extracts were centrifuged at maximum speed for 10 min at 4°C. Supernatant was mixed with SDS loading buffer and boiled for 10 min. Protein extracts were separated by SDS-PAGE, transferred to PVDF membranes, and blocked for 1 hour. Membranes were probed with mouse monoclonal anti-Actin (A0480, Sigma-Aldrich) and mouse monoclonal anti-FLAG (F1804, Sigma-Aldrich), followed by HRP-conjugated anti-mouse secondary antibody (1706516, Bio-Rad). For detection of biotinylated proteins, membranes were blocked with 2% BSA for 1 hour and incubated with Streptavidin-HRP (S911, Thermo Fisher Scientific). After washing, signals were visualized using the SuperSignal West Dura chemiluminescent substrate (Thermo Fisher Scientific).

### Fluorescence imaging and Photobody analysis

Fluorescence microscopy was performed using a Zeiss Axio Observer 7 inverted microscope equipped using either EC Plan-Neofluar 40×/0.8, or C-Apochromat 63×/1.2 objectives. Images were obtained using a Colibri 7 LED light source and standard Zeiss filter sets: DAPI (Zeiss filter set 49, excitation 385/30 nm, emission 445/50 nm), CFP (Zeiss filter set 47, excitation 423/44 nm, emission 480/40 nm), and YFP (Zeiss filter set 46, excitation 511/44 nm, emission 535/30 nm). Image acquisition was carried out with ZEN Blue software (Zeiss). Volume and number of photobodies were analyzed using the object analyzer tool of Huygens Essential Software (Scientific Volume Imaging) (Van Buskirk *et al*., 2014; Yoo *et al*., 2019b).

### Proximity labeling and sample preparation for LC-MS/MS

Proximity labeling was performed as described previously with minor modifications (Mair *et al*., 2019). Seedlings expressing miniTurbo fusions were grown for 3 days in darkness, exposed to 36 μmol m^−2^ s^−1^ red light for 24 hours, and treated with 50 μM biotin for 3 hour. Approximately 1 g of tissue per replicate was harvested, frozen in liquid nitrogen, and homogenized in extraction buffer (50 mM Tris-HCl pH7.5, 150 mM NaCl, 0.1% SDS, 1% Triton X-100, 0.5% sodium deoxycholate, 1mM EGTA, 1mM DTT, protease inhibitor cocktail, 1 mM PMSF, 40 μM MG132, 40 μM MG115). Free biotin was removed by Cytiva PD-10 desalting columns (Thermo Fisher Scientific). Biotinylated proteins were captured with Dynabeads MyOne Streptavidin beads (Thermo Fisher Scientific), washed with a series of buffers, and resuspended with equilibration buffer (50mM Tris-HCl pH7.5, 150 mM NaCl, 0.1% SDS, 1% Triton X-100, 0.5% sodium deoxycholate, 1mM EGTA, 1mM DTT). Biotinylated proteins on beads were taken to the Mass Spectrometry and Proteomics Core Facility at the University of Utah.

### LC-MS/MS analysis and data processing

Streptavidin-conjugated beads containing biotinylated proteins from PBmTb and mTb control lines were digested on bead with trypsin. Peptides were reduced, alkylated, desalted using Pierce Peptide Desalting Columns (Thermo Fisher Scientific), and then resuspended in 0.1% formic acid for analysis.

Peptide separation was performed on a nanoElute 2 system (Bruker Daltonics) coupled to a timsTOF Pro2 mass spectrometer equipped with a nanoelectrospray source. Peptides were analyzed by reversed-phase nano-LC-MS/MS using a C18 column and a 45-min gradient of 0.1% formic acid in water (buffer A) and acetonitrile (buffer B). Data were acquired in PASEF data-dependent acquisition (DDA) MS/MS scan mode and processed at the the Mass Spectrometry and Proteomics Core Facility at the University of Utah.

Raw data were processed using FragPipe v20.0 software against *Arabidopsis thaliana* UniProt reference proteome. Protein abundances based on peak intensity were quantified using DIA-NN software v1.8.1. Variable modifications included methionine oxidation and cysteine carbamidomethylation, with mass tolerances of 15 ppm for precursor and 10 ppm for fragment ions. A false discovery rate of 1% and 5% was applied at both protein and peptide levels, and proteins were required to have at least one unique peptide.

## ACKNOWLEDGMENTS

This work was supported by start-up funds from the University of Utah (C.Y.Y) and by National Institutes of Health (NIH) Training Program in Developmental Biology T32HD007491 (K.O.). Proteomics mass spectrometry analysis was performed at the Mass Spectrometry and Proteomics Core Facility at the University of Utah. We thank Dominique Bergman for sharing and depositing the miniTurbo-YFP-NLS construct to Addgene (#127370).

## AUTHOR CONTRIBUTIONS

K.O. and C.Y.Y. designed research; K.O., J.-H.L, M.D., T.M.D., A.M.M., and C.Y.Y. performed research; K.O., A.M.M., and C.Y.Y. analyzed data; K.O. and C.Y.Y. wrote the paper.

---

**Figure S1.** phyB-miniTurbo does not affect phyA-mediated far-red light signaling. (a) Representative images of 4-day-old Col-0, *phyA-211/phyB-9* (*phyA/phyB*), *PBmTb #3-1, PBmTb #10-1*, and *mTb #2-32* seedlings grown in 10 μmol m^-2^ s^-1^ FR light. (b) Box-and-whisker plots showing hypocotyl lengths of seedlings shown in (a). Boxes indicate the 25^th^ to 75^th^ percentiles with median values shown as horizontal lines; whisker extends to the minimum and maximum values. Different letters represent significant differences (one-way ANOVA with Tukey’s HSD test, *P* < 0.01).

**Figure S2.** Morphological and transcriptional characterization of PBmTb seedlings during de-etiolation. (a) Representative images of PBmTb #3-1 seedlings transferred from darkness to light for the indicated durations (8, 16, and 24 hour) and under continuous red light (R_C_). (b) Transcript levels of plastid-encoded photosynthetic genes *psbA* and *rbcL* relative to *PP2A*. Data represent means ± SD from three biological replicates. Different letters indicate statistically significant differences (one-way ANOVA with Tukey’s HSD test, *P* < 0.01).

**Figure S3.** Quality assessment and phyB enrichment in the PBmTb proximity proteomics dataset. (a) Principal component analysis (PCA) of proteomic datasets from PBmTb #3-1 and mTb control seedlings showing distinct clustering by genotype across three biological replicates. (b) Abundance of phyB (UNIPROT ID: P14713) detected in PBmTb and mTb samples before and after normalization. Detection of phyB reflects self-biotinylation of the PBmTb fusion protein, confirming active miniTurbo ligase and efficient recovery of the bait protein.

